# Transcriptomic profiling of high-density Giardia foci encysting in the murine proximal intestine

**DOI:** 10.1101/114983

**Authors:** JK Pham, C Nosala, EY Scott, KF Nguyen, KD Hagen, HN Starcevich, SC Dawson

## Abstract

*Giardia* is a highly prevalent, understudied protistan parasite causing significant diarrheal disease worldwide. Its life cycle consists of two stages: infectious cysts ingested from contaminated food or water sources, and motile trophozoites that colonize and attach to the gut epithelium, later encysting to form new cysts that are excreted into the environment. Current understanding of parasite physiology in the host is largely inferred from transcriptomic studies using *Giardia* grown axenically or in co-culture with mammalian cell lines. The dearth of information about the diversity of host-parasite interactions occurring within distinct regions of the gastrointestinal tract has been exacerbated by a lack of methods to directly and non-invasively interrogate disease progression and parasite physiology in live animal hosts. By visualizing *Giardia* infections in the mouse gastrointestinal tract using bioluminescent imaging (BLI) of tagged parasites, we recently showed that parasites colonize the gut in high-density foci that may cause localized pathology to the epithelium. Encystation is also initiated in these foci throughout the entire course of infection, yet how the physiology of parasites within high-density foci in the host gut differs from that of cells in laboratory culture is unclear. Here we use BLI to precisely select parasite samples from high-density foci in the proximal intestine to interrogate *in vivo Giardia* gene expression in the host. Relative to axenic culture, we noted significantly higher expression (> 10-fold) of oxidative stress, membrane transporter, and metabolic and structural genes associated with encystation in the high-density foci. These differences in gene expression within parasite foci in the host may reflect physiological changes associated with high-density growth in localized regions of the gut. We also identified and verified six novel cyst-specific proteins, including new components of the cyst wall that were highly expressed in these foci. Our *in vivo* transcriptome data support an emerging view that parasites encyst early in localized regions in the gut, possibly as a consequence of nutrient limitation, and also impact local metabolism and physiology.

## Introduction

*Giardia lamblia* is a zoonotic protozoan parasite causing acute and chronic diarrheal disease in over one billion people worldwide, primarily in areas lacking adequate sanitation and water treatment [1; 2]. Giardiasis is a serious disease of children, who may experience substantial morbidity including diarrhea, malnutrition, wasting, and developmental delay [3; 4; 5]. In the United States, giardiasis is the most frequently diagnosed cause of water-borne diarrheal disease, and commonly affects travelers and immunosuppressed individuals [6]. The estimated failure rates of up to 20% for standard treatments [7] and reports of drug resistance [8; 9; 10] underscore the need for new therapeutic treatments for this widespread and neglected diarrheal disease.

Giardiasis may be either acute and/or chronic, and infection is generally accompanied by abdominal cramps, gas, nausea, and weight loss. Giardiasis may also result in a severe form of malabsorptive diarrhea presenting as a fatty, watery stool [11]. Trophozoites are not invasive and giardiasis does not produce a florid inflammatory response; however, giardiasis is associated with villus shortening, enterocyte apoptosis, hypermobility, and intestinal barrier dysfunction [12]. The mechanisms by which *Giardia* colonization of the gastrointestinal tract induces diarrheal disease remain unclear.

*Giardia* differentiates into two morphological forms during its life cycle: a motile, multi-flagellated trophozoite that colonizes the host small intestine; and an infective cyst that is shed into the environment [13]. Following ingestion by a mammalian host, cysts transform into motile trophozoites as they pass into the gastrointestinal tract. Trophozoites navigate the lumen of the small intestine and attach to the microvilli of the small intestine via the ventral disc, yet do not invade the epithelium [6]. Attachment to the gut epithelia allows the parasite to resist peristaltic flow and proliferate in this low oxygen, nutrient rich environment. Trophozoites are eventually triggered to differentiate into cysts, which exit the host and are disseminated with feces into the environment.

The lack of accessibility to the gastrointestinal tract [14; 15; 16] has limited our understanding of *in situ Giardia* physiology in the host. Parasite physiology and metabolism have been inferred from axenic laboratory culture in a non-defined medium [17] or from co-culture with intestinal epithelial cell lines [14; 16; 18]. However, co-incubation studies of trophozoites with intestinal cell lines may not accurately reflect *in vivo* parasite physiology. *Giardia* infects many mammalian hosts, and animal models of giardiasis include adult [19; 20] or suckling mice [21] or adult gerbils [22] infected with either *Giardia lamblia or G. muris* isolates [23]. While the adult mouse model of giardiasis is commonly used to evaluate anti-giardial drugs [24], previous *in vivo* parasite studies lack precision due to tissue sampling throughout the intestinal tracts of infected animals without corresponding knowledge of *Giardia* colonization.

Our recent development of methods for non-invasive imaging of bioluminescent *Giardia* parasites permits unprecedented access to real-time host-parasite interactions, allowing us to revise decades-old assumptions about *Giardia* infection and encystation dynamics in living hosts. BLI enables sensitive quantification and live reporting of transcriptional activity of promoter-luciferase fusions [25; 26; 27] but also protein-luciferase fusions [28]. BLI is used commonly to monitor parasitic infection dynamics during malaria, leishmaniasis, trypanosomiasis, and toxoplasmosis [29; 30; 31], as well as bacterial colonization of the intestine [32]. The use of imaging methods to evaluate *in vivo* giardiasis has allowed the precise longitudinal and spatial monitoring of the dynamics of infection, and provides an improved method to evaluate anti-giardial drugs in a relevant animal model of giardiasis.

Using BLI, we recently demonstrated that metabolically active *Giardia* primarily colonizes the proximal small intestine, rather than the mid-jejunum as has been previously reported [15; 19; 33; 34]. By imaging mice infected with *Giardia* expressing firefly luciferase under the control of encystation-specific [32; 35; 36; 37; 38] promoters, we determined that encystation is initiated early in the course of infection and is correlated to dense sites, or foci. These foci persist in the gastrointestinal tract throughout the course of infection.

Here we interrogate the *in vivo* physiology of high-density *Giardia* foci in the proximal small intestine and compare this to physiology in axenic culture. We observe that also upregulated in parasite metabolism in the foci is shifted toward encystation, and that genes involved in response to oxidative stress are also upregulated. Furthermore, we observe that trophozoites in high-density foci are encysting and exhibit a expression profiles to *in vitro Giardia* cells in profile similar to the profile seen during mid-to-late encystation (7-22 hours in encystation media) [39]. Finally, we identify and confirm six new encystation genes that are highly expressed in the foci.

## Materials and Methods

### Giardia trophozoite and encystation culture conditions

The *G. lamblia* (ATCC 50803) bioluminescent reporter *P_GDH_-Fluc* [40] strain, all C-terminal GFP-fusion strains, and the wild type WBC6 strain were cultured in modified TYDK medium (also known as TYI-S-33 or Keister’s medium [17]) supplemented with bovine bile and 5% adult and 5% fetal bovine serum [56] in sterile 16 ml screw-capped disposable tubes (BD Falcon), and incubated upright at 37°C without shaking. Encystation was induced *in vitro* by decanting TYDK medium from 24-hour cultures (roughly 30% confluent) and replacing with encystation medium modified by the addition of 0.5 grams/liter bovine bile, pH 7.8 [39]. After 24 hours, cysts settled and were harvested for subsequent imaging analyses.

### *In vivo* and *ex vivo* bioluminescence imaging of mice with spatial sampling of *Giardia* colonization

Animal studies were performed with IACUC approval at the University of California, Davis (Scott C. Dawson, PI). Protocols for *in vivo and ex vivo* bioluminescence imaging of *Giardia-infected* mice were as previously described [40]. Specifically, four eight week old, female C57/B6/J mice (Jackson Laboratories) were maintained on *ad libitum* water and alfalfa-free irradiated rodent pellets (Teklad 2918). Water was supplemented with 1 mg/ml ampicillin and neomycin (Teknova) for five days prior to infection [41] to promote parasite colonization, and mice were maintained on antibiotics for the entire experiment. For infections, each animal was gavaged with 1 × 10^7^ G. *lamblia P_GDH_-Fluc* strain trophozoites in 100 μL phosphate-buffered saline [40].

For non-invasive monitoring of parasite colonization using *in vivo* BLI, mice were sedated, D-luciferin (30 mg/kg) was then injected intraperitoneally, and bioluminescence was imaged using an IVIS Spectrum (PerkinElmer) with an open emission filter [40]. Photons were quantified using an ultra-sensitive CCD camera (IVIS Spectrum) and the resulting heat maps of bioluminescent photon emission intensity were overlaid on still images of anaesthetized animals. To allow the D-luciferin to distribute throughout the body, images were collected with two-minute exposures constantly over 8-10 minutes until the bioluminescent signal stabilized. The exposure time for final image collection ranged from two to five minutes due to differences in the signal strength. Bioluminescence was quantified using LivingImage software to draw a region of interest (ROI) around each mouse, from front paws to anus. BLI data was quantified as total flux (photons/second) for exposure time-independent quantification of signal intensity.

To image the *ex vivo* spatial location and density of *Giardia* in murine gastrointestinal tract, sedated mice were euthanized on day 3 or day 7 post-gavage by cervical dislocation. The entire gastrointestinal tract was quickly dissected from esophagus to anus and positioned within a plastic Petri dish for *ex vivo* imaging using thirty second exposures [40]. ROI analysis was used to quantify bioluminescence (LivingImage). Total gastrointestinal tract signal was analyzed within a circle encompassing the entirety of the petri dish. The stomach, proximal SI (first half), distal SI (second half), cecum, and large intestine were traced using the free-hand tool and total flux was recorded as stated above (Supplemental Table 1). Spatial imaging of bioluminescence in the entire gastrointestinal tract specified the particular intestinal segments enriched for high-density *Giardia* foci of colonization. Small intestinal segments with such foci were excised for downstream RNA extraction and stored in RNAlater (Life Technologies) at -80°C, or alternatively, were fixed in 10% phosphate-buffered formalin for histology.

Histological sections of these intestinal samples were stained and visualized using light microscopy, confirming the presence of *Giardia* trophozoites in selected intestinal samples [40]. Protocols for preparation of histological slides were the same as those described in Chen et al. [42] and were performed by the UC Davis Anatomic Pathology and IHC Laboratory (Davis, CA). Briefly, a subset of the excised bioluminescent intestinal samples was fixed in 10% phosphate-buffered formalin for 24 hours. Specimens were dehydrated in a series of graded ethanol (70%, 96% and 99%) and embedded in paraffin. Five-micron sections were cut perpendicular to the mucosa surface and the paraffin was cleared from the slides with coconut oil (over 15 min, 60°C). Sections were rehydrated in 99%, 96% and 70% ethanol followed by a 10 minute wash in water and stained with hematoxylin and eosin (HE). All HE slides were visualized via brightfield microscopy using a Leica DMI 6000 wide-field inverted fluorescence microscope with PlanApo 10X and 40X objectives.

### Total RNA extraction and RNA-seq of gastrointestinal samples in infected animals

Total RNA was extracted by from *Giardia-infected* intestinal segments by resuspending the tissue in RNA STAT 60 solution (Tel-Test, Inc.) and repeatedly pipetting through a glass pipette until tissue homogenization was complete. All samples were kept on ice during tissue homogenization. Total RNA was purified by phenol-chloroform extraction and isopropanol precipitation. Poly(A) selection was performed with one microgram of total RNA. Illumina sequencing libraries were constructed and ribosomal RNA was depleted (UC Davis DNA Technologies Core Facility, Davis, CA) for downstream Illumina sequencing. 120-180 million 50 bp, single end reads were subsequently generated via Illumina sequencing (HiSeq 2500 system).

### Trimming and processing of transcriptome sequences

Raw sequences were trimmed using the sliding window trimmer Sickle [43]. Two pipelines were used for mapping and transcript quantification (Figure 1D). First, the reads were mapped to the *Giardia* assemblage A genome (WBC6, release 5.0) using TopHat (version 2.0.14) [44]. The TopHat output BAM files were processed and assembled into transcripts by Cufflinks (version 2.2.1) [45] and normalized to fragments per kilobase of transcript per million fragments mapped (FPKM, see Supplemental Table 2). Differentially expressed genes were identified using the cuffdiff function of the Cufflinks package (version 2.2.1) [46]. Differentially expressed genes were defined using a threshold cutoff false-discovery rate (FDR) of 5%. The R package cummeRbund was used to generate summary figures for the resulting RNA-seq data, such as heat maps, PCA plots, and gene clusters of differentially expressed genes (via partitioning tools) [46]. The second pipeline mapped the reads back to the *Giardia* transcriptome (GiardiaDB-5.0_GintestinalisAssemblageA.gff) with Kallisto [47], using the recommended 30 bootstraps per sample. The companion package, Sleuth [47], was used for differential gene expression analysis, using the Wald test and significance q-value cutoff of 0.05. Transcripts showing concordant significant differential gene expression between Cuffdiff and Sleuth statistical packages were selected as confident DE genes.

**Figure 1.**
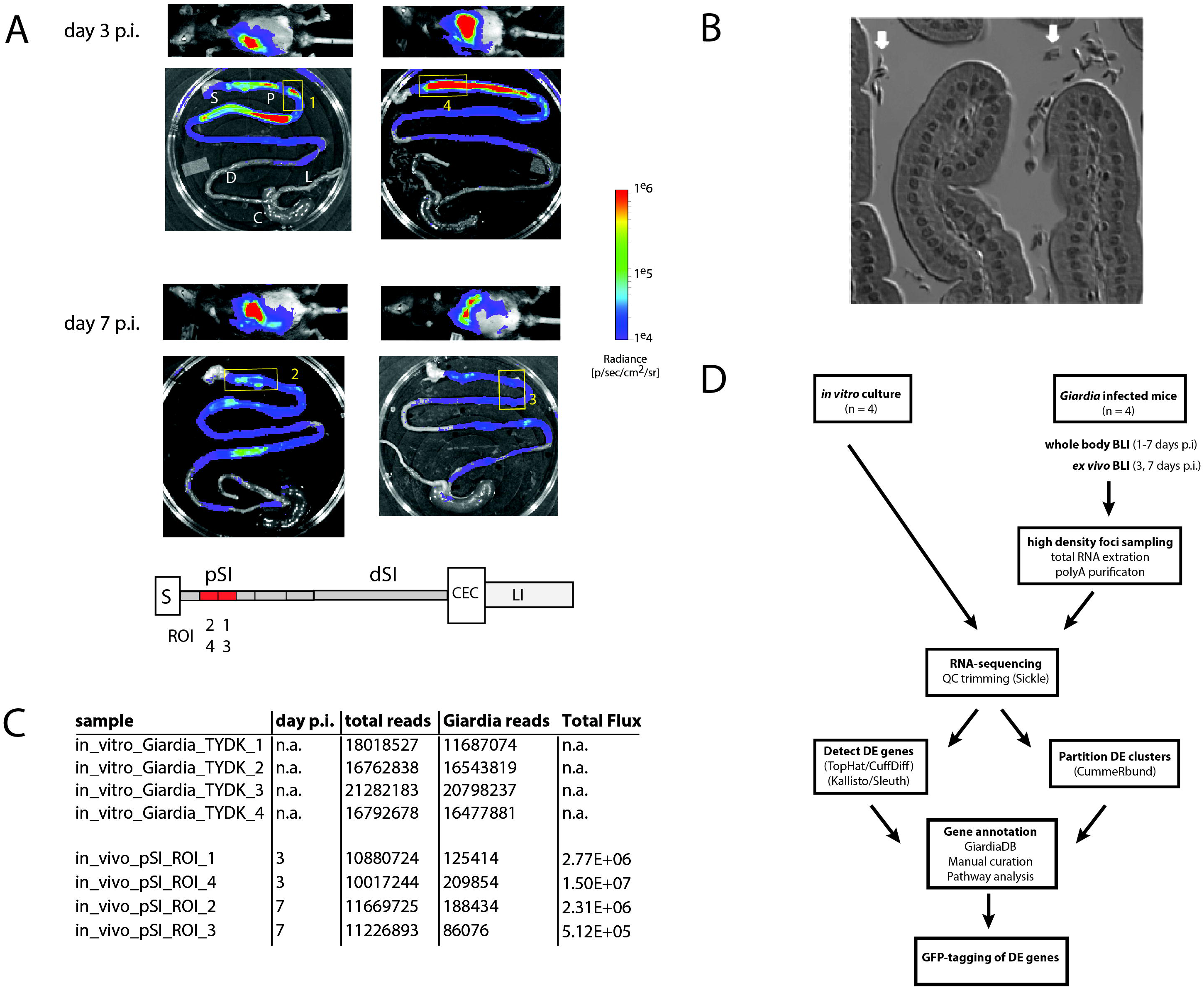
Bioluminescent imaging allows precise sampling of Giardia foci in the proximal small intestine for transcriptomic analyses. A cohort of four mice were infected with one million trophozoites by gavage of the *P_GDH_-FLUC Giardia* strain [40]. Whole body images were non-invasively collected for parasite bioluminescence from the constitutive *P_GDH_-FLUC* bioreporter (A), and mice were subsequently sacrificed at 3 or 7 days post infection (p.i.). Image overlays display the *P_GDH_-FLUC* bioluminescence intensity, with the highest signal intensity shown in red and the lowest in blue. The colored radiance scale indicates the photon flux (p/s/cm^2^/sr) for each intestinal segment. *Ex vivo* imaging defined four foci in the proximal small intestine (yellow numbered boxed regions) subsequently excised for transcriptomic sequencing and analysis (A). In the schematic, regions of the gastrointestinal tract are marked as: S = stomach, pSI = proximal small intestine, dSI = distal small intestine, CEC = cecum, and LI = large intestine, with the positions of the *Giardia* foci (ROI1-RO14) within the excised fragments in red. In B, the enrichment of *Giardia* trophozoites (white arrow) in a representative bioluminescent small intestinal sample was verified using light microscopy of histological sections. Panel C summarizes the total transcriptomic reads for each *in vitro* culture replicate (TYDK) and for each transcriptomic sample (ROI1-ROI4) from the proximal small intestine (pSI). The total reads include those from the host (mouse) before the *Giardia* reads were computationally mapped the *Giardia* genome. The total flux (p/s/cm^2^/sr) was quantified for that particular sample. The overall transcriptomic analysis is summarized in panel D.

For visualization, FPKM values of the 475 confident DE genes were hierarchically clustered. Biclustering was performed by Pearson correlation for transcript expression and Spearman correlation of expression profiles associated with each sample. The heatmap (Supplemental Table 3) was plotted using the heatmap.2 function of “gplots” the R software package (https://CRAN.R-project.org/package=gplots).

### Functional transcriptome annotation, metabolic pathway mapping, and partitioning analyses

Cellular functions were inferred for significant differentially expressed *Giardia* genes using analytical tools associated with the GiardiaDB (http://giardiadb.org/giardiadb/) [48], with subsequent manual inspection and curation of homology, functional domains, and cellular process designations according to prior experimental and bioinformatic evidence. GiardiaDB analytical tools were used to compare *in vivo* transcriptomic reads to prior transcriptomic and proteomic experimental studies [34; 49; 50]. In addition, the cummeRbund R package was used for partitioning analysis to predict nine differentially expressed gene clusters using the Jenson-Shannon distance binning methods with the cummeRbund software package [46], and compared to both the *in vivo* foci gene clusters and prior Giardia encystation and Caco-2-associated transcriptome studies [34; 49; 50].

### C-terminal GFP fusion strain construction of select differentially expressed genes

To confirm the cellular localization of novel differentially expressed genes identified in the *in vivo* transcriptome, 15 differentially expressed *Giardia* genes (GL50803_7797, GL50803_4705, GL50803_5258, GL50803_7710, GL50803_15499, GL50803_10822, GL50803_15427, GL50803_24412, GL50803_14748, GL50803_6679, GL50803_88960, GL50803_5515, GL50803_14567, GL50803_14926, GL50803_9354) were GFP-tagged via our laboratory’s Gateway cloning pipeline [51]. We also tagged fourteen genes that were more highly expressed in *in vitro* culture. The C-terminal GFP fusion constructs included approximately 200–250 nucleotides upstream of the gene, the gene itself in frame with GFP, and a puromycin resistance cassette [51]. The *Giardia* strain WBC6 was electroporated with 20 μg of GFP-fusion plasmids, and transformed strains were maintained under antibiotic selection (50 μg/ml puromycin) for at least two weeks [51].

### Immunostaining and imaging of encysting GFP-tagged strains

To visualize encystation vesicles and the mature cyst, GFP-tagged trophozoites were encysted using established *in vitro* encystation methods [52]. Sterile deionized water was then added to all experimental culture tubes to lyse incompletely encysted cells. Water resistant cysts were stored at 4°C for subsequent imaging. Cyst wall protein 1 (CWP1) was visualized by immunostaining fixed encysting trophozoites attached to coverslips using a 1:200 dilution of anti-CWP1 primary antibody (Waterborne, Inc., New Orleans, LA) and a goat anti-mouse antibody conjugated to Alexa Fluor 594 (Invitrogen) as previously described [52]. Coverslips were mounted onto slides using Prolong Gold Antifade Solution with DAPI (Invitrogen).

Three dimensional stacks of immunostained samples were acquired using the Metamorph image acquisition software (MDS Technologies) with a Leica DMI 6000 wide-field inverted fluorescence microscope with a PlanApo 100X, NA 1.40 oil immersion objective. Serial sections of GFP-tagged strains were acquired at 0.2 μm intervals and deconvolved using Huygens Professional deconvolution software (SVI). Two dimensional maximum intensity projections were created from the 3D stacks for presentation.

## Results

### Bioluminescent imaging allows precise sampling of regions of the proximal small intestine with high-density Giardia foci for transcriptomics

A cohort of four of mice were inoculated with the constitutive metabolic bioreporter strain *P_GDH_-Fluc* [40]. *Giardia-specific* bioluminescence was quantified non-invasively in animals from one to seven days post-inoculation (Figure 1A and Methods). Two animals were sacrificed at day 3 p.i., and two more at day 7 p.i. *Ex vivo* imaging of the gastrointestinal tracts revealed dense foci of bioluminescent signal primarily in the proximal and distal small intestine (Figure 1A); bioluminescent flux directly correlates with the number of active *Giardia* parasites [40]. Proximal intestinal segments from each gastrointestinal tract were sampled, and the bioluminescent flux of the *Giardia* foci (5.12 × 10^5^ to 1.5 × 10^7^ photons/second/cm (Figure 1B)) was determined. Representative proximal small intestinal samples (ROI) from each gastrointestinal tract were selected for transcriptomic analyses. The histology of these segments indicated no verifiable host tissue damage (Methods), but did indicate obvious colonization of these regions by *Giardia* trophozoites (Figure 1A and 1C).

Sufficient purified total RNA was obtained from four *in vivo* intestinal samples and was used to create RNA-seq libraries for sequencing (Methods and Figure 1B). Four additional RNA-seq libraries were created and sequenced from four independent *in vitro Giardia P_GDH_-Fluc* cultures grown axenically in standard medium (Methods). Raw sequence reads generated for the RNA-seq samples ranged from 10.9 to 21.3 million (Figure 1B). The majority of RNA-seq reads from the *in vitro* culture (TYDK) datasets (97.7% to 98.7%) were mapped back to the WBC6 genome. Due to the high proportion of host total RNA in the *ex vivo* samples, fewer RNA-seq reads (0.8% to 1.9%) were mapped to the WBC6 genome; however, these numbers were comparable between individual *in vivo* samples and were sufficient for subsequent analyses of differential gene expression (Figure 1B and Methods).

### Identification of 475 differentially expressed (DE) genes between in vitro culture and in vivo foci transcriptomes

Transcriptomic comparisons and analyses between *in vitro* and *in vivo* samples were performed using the TopHat, Cufflinks, CummeRbund, Kallisto, and Sleuth packages as outlined in the bioinformatic strategy in Figure 1D and [46]. A comparison of the normalized read (FPKM) profiles of the *in vitro* and *in vivo* transcriptomes indicate that the four *in vitro* profiles were very similar to one another, as were the four *in vivo* foci profiles (see heat map, Figure 2A and Supplemental Table 2). Significant differences in gene expression profiles between the *in vitro* and *in vivo* samples (Figure 2A) were quantified and summarized using two different computational methods (Cuffdiff and Sleuth). Cuffdiff and Sleuth identified 1073 and 1336 differentially expressed genes, respectively, between the *in vitro* and *in vivo* datasets. Four hundred seventy-five differentially expressed genes were concordant between the two methods (Figure 2B). One hundred eighty-seven of the concordant differentially expressed genes had increased expression in the *in vivo* transcriptomes relative to the *in vitro* transcriptomes (Figure 2C).

**Figure 2.**
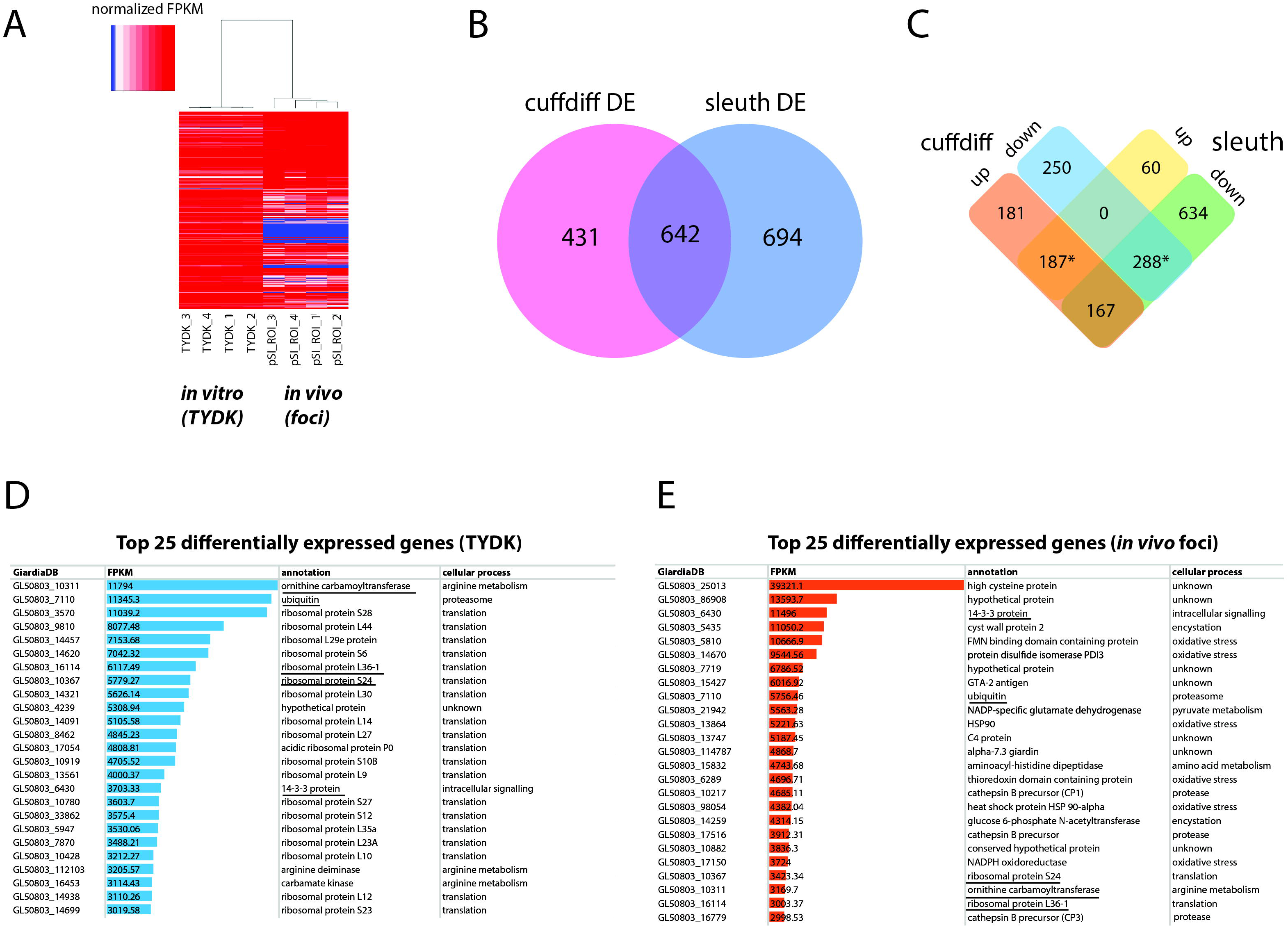
Summary of differentially expressed *in vitro* and *in vivo* foci transcriptomes using Cuffdiff and Sleuth. Panel A depicts a heat map of normalized FPKM comparing RNA-seq datasets from the *Giardia* grown *in vitro* culture (TYDK) to *Giardia* transcriptomes associated with the *in vivo* foci excised from the proximal small intestine (pSI) (see Figure 1). Two computational methods of identifying differentially expressed transcripts in the *in vitro* (TYDK) and *in vivo* (foci) datasets have some overlap (B). Overlap in both upregulated and downregulated genes in the *in vivo* transcriptomes was calculated and shown in (C). The asterisk indicates upregulated genes and downregulated genes that were identified using both methods. The top 25 expressed genes (FPKM) present in the *in vitro* (D) and *in vivo* foci transcriptomes (E) are also ranked. Underlined gene names are those that occur in the top 25 expressed genes in both the *in vitro* and the *in vivo* transcriptome datasets.

Two hundred eighty-eight of the concordant differentially expressed genes had decreased expression in the *in vivo* foci transcriptomes relative to *in vitro* transcriptomes. All subsequent analyses focused on the concordant genes with expression levels that varied more than two-fold between the *in vivo* and *in vitro* transcriptomes. These included 187 genes that were more highly expressed *in vivo* foci and 166 that were more highly expressed *in vitro* culture. (Figure 2C and Supplemental Tables 4 and 5). Sixty-one genes that were expressed *in vitro* were not expressed in the *in vivo* foci transcriptomes (Supplemental Table 6).

Statistically significant increases in gene expression ranging from 2.5-to 153-fold were observed for *Giardia* in *in vivo* foci relative to trophozoites cultured *in vitro.* With respect to highly expressed genes, transcripts associated with translation and arginine metabolism were the most abundant in the *in vitro* transcriptomes (Figure 2D and Supplemental Tables 4 and 6). In contrast, transcripts associated with encystation, oxidative stress, or unknown functions are among the most abundant in the *in vivo* foci transcriptomes (Figure 2E). Despite significant differences between *in vivo* and *in vitro* expression profiles, five genes (ornithine carbamoyl transferase (OCT, 10311), ubiquitin (7110), 14-3-3 protein (6430), ribosomal protein L36-1 (16114) and ribosomal protein S24 (10367) are among the top transcribed genes in both transcriptome datasets (Figure 2D and 2E).

### Encystation-specific and oxidative stress genes are significantly upregulated in the in vivo foci

Of the 475 differentially expressed genes, 25 genes were expressed more than 10-fold higher in the *in vivo* foci relative to cultured cells; however, only 8 genes were expressed at levels more than 10-fold higher in cultured cells than *in vivo* (Figure 3A and Supplemental Table 4 and 5). The expression of 85 genes was elevated more than 5-fold in foci relative to cultured trophozoites, and there were 29 genes whose expression was more than 5-fold higher in the *in vitro* transcriptomes relative to the *in vivo* transcriptomes (Supplemental Tables 4 and 5).

**Figure 3.**
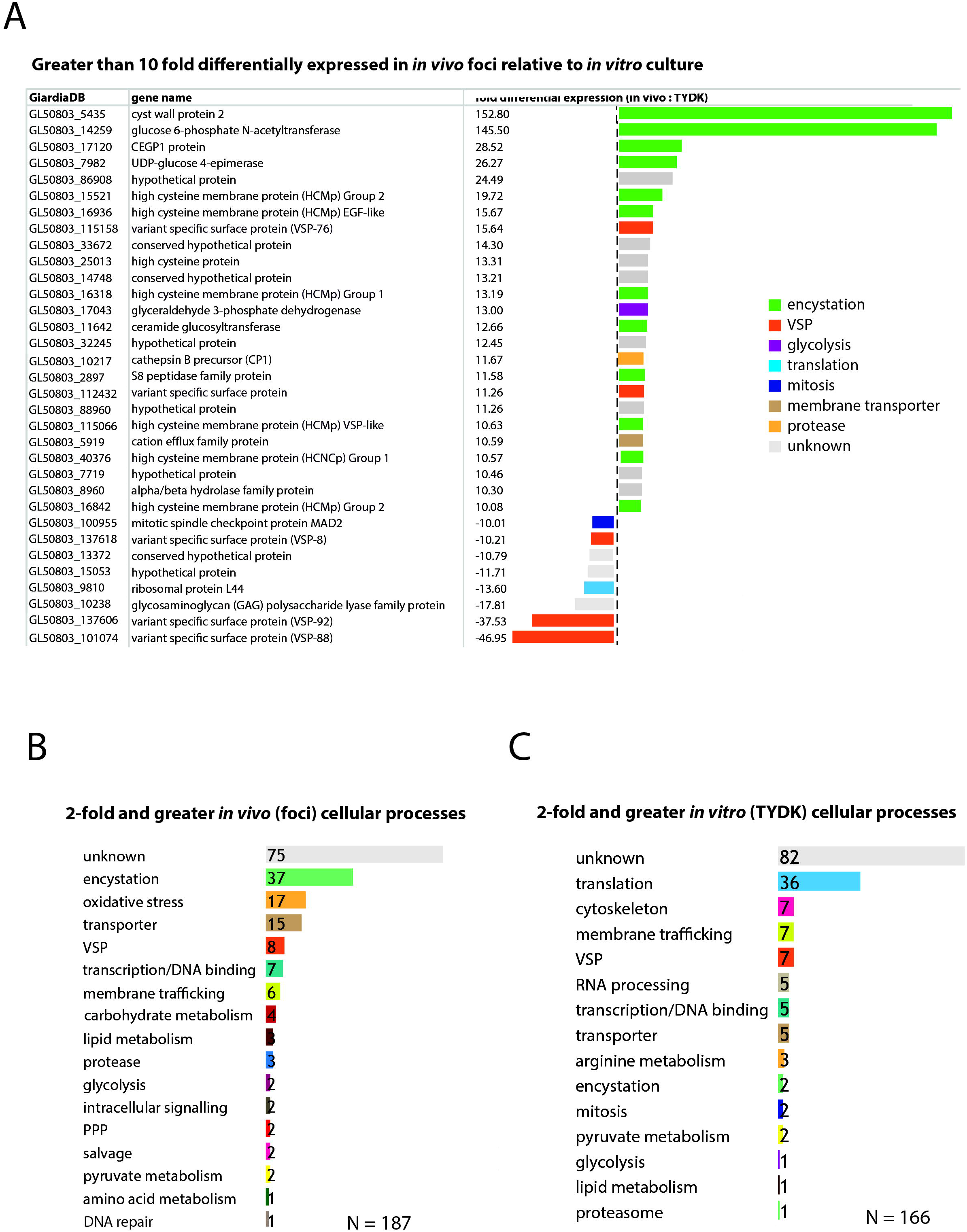
Encystation and oxidative stress genes are the top differentially expressed genes associated with the *in vivo* foci transcriptomes. In panel A, genes that are greater than 10-fold differentially expressed in the *in vivo* foci transcriptomes relative to *in vitro* culture are ranked and colored according to cellular process (e.g., encystation). Genes differentially expressed greater than two-fold are ranked according to the number that are associated with a particular cellular process in the *in vivo* foci transcriptome (B) and the *in vitro* (TYDK) transcriptome (C).

In contrast to the genomes of model organisms, many genes within the WBC6 genome lack functional gene annotations associated with *Giardia-specific* processes (e.g., encystation or pyruvate metabolism). Thus, we manually curated the functional annotations of genes whose expression was more than 2-fold higher *in vivo* (187 genes) or *in vitro* (166 genes) and summarized the functional categories associated with these statistically significant differentially expressed genes (Figure 3). Ten of the 25 genes with more than 10-fold higher expression in the *in vivo* foci are associated with encystation-specific processes. These genes encode components of the cyst wall (CWP2, 153X higher *in vivo*), enzymes in the UDP-N-acetylgalactosamine biosynthetic pathway (GL50803_14259, 146X higher *in vivo),* and high cysteine membrane proteins (HCMPs, 10X-20X higher *in vivo*). Eight genes with more than 10-fold higher expression in the *in vivo* foci have unknown or uncharacterized functions (Figure 3A and Supplemental Table 4).

Encystation-specific gene expression in *in vivo* foci becomes even more apparent when we consider the functional annotations of genes with more than 2-fold higher expression in *in vivo* foci relative to *in vitro* culture are summarized. Thirty-seven differentially expressed genes are associated with encystation, 17 with oxidative stress responses, and 15 with membrane transporter functions (Figure 3B and Supplemental Table 4). This is in contrast to genes highly expressed *in vitro* culture, which include cytoskeletal genes and other genes associated with housekeeping functions such as translation, RNA processing, and transcription (Figure 3C and Supplemental Table 5). Different types of genes with membrane trafficking functions are increased in the two transcriptome dataset. About equal numbers of unique variant specific surface proteins (or VSPs, [53]) are differentially expressed amongst the *in vitro* (7 VSPs) and *in vivo foci* (8 VSPs) transcriptomes.

Lastly, genes with unknown or uncharacterized function are the most abundant category in both the *in vivo* foci transcriptome (75/187) and the *in vitro* transcriptome (82/166) datasets.

### *In vivo* foci display increased expression of genes associated with the UDP-N-acetylgalactosamine, pentose phosphate, and glycolytic and pyruvate pathways

To evaluate the degree to which the differential gene expression we observed is associated with specific metabolic pathways in *Giardia,* we mapped metabolic genes to four catabolic and biosynthetic pathways (Figure 4) using manual curation, as well as GO term and metabolic pathway enrichment tools available from GiardiaDB [48]. Specifically, we mapped the transcriptional upregulation of genes associated with *in vivo* foci to glycolysis and pyruvate metabolism, the pentose phosphate pathway, the UDP-N-acetylgalactosamine pathway, and the arginine dihydrolase pathway (Figure 4). First, we noted that in *in vivo* foci, the three key enzymes of the arginine dihydrolase pathway (OCT, ADI, CK) showed 3.7-6.4X less gene expression in *in vivo* foci than in trophozoites grown in *in vitro* culture.

**Figure 4.**
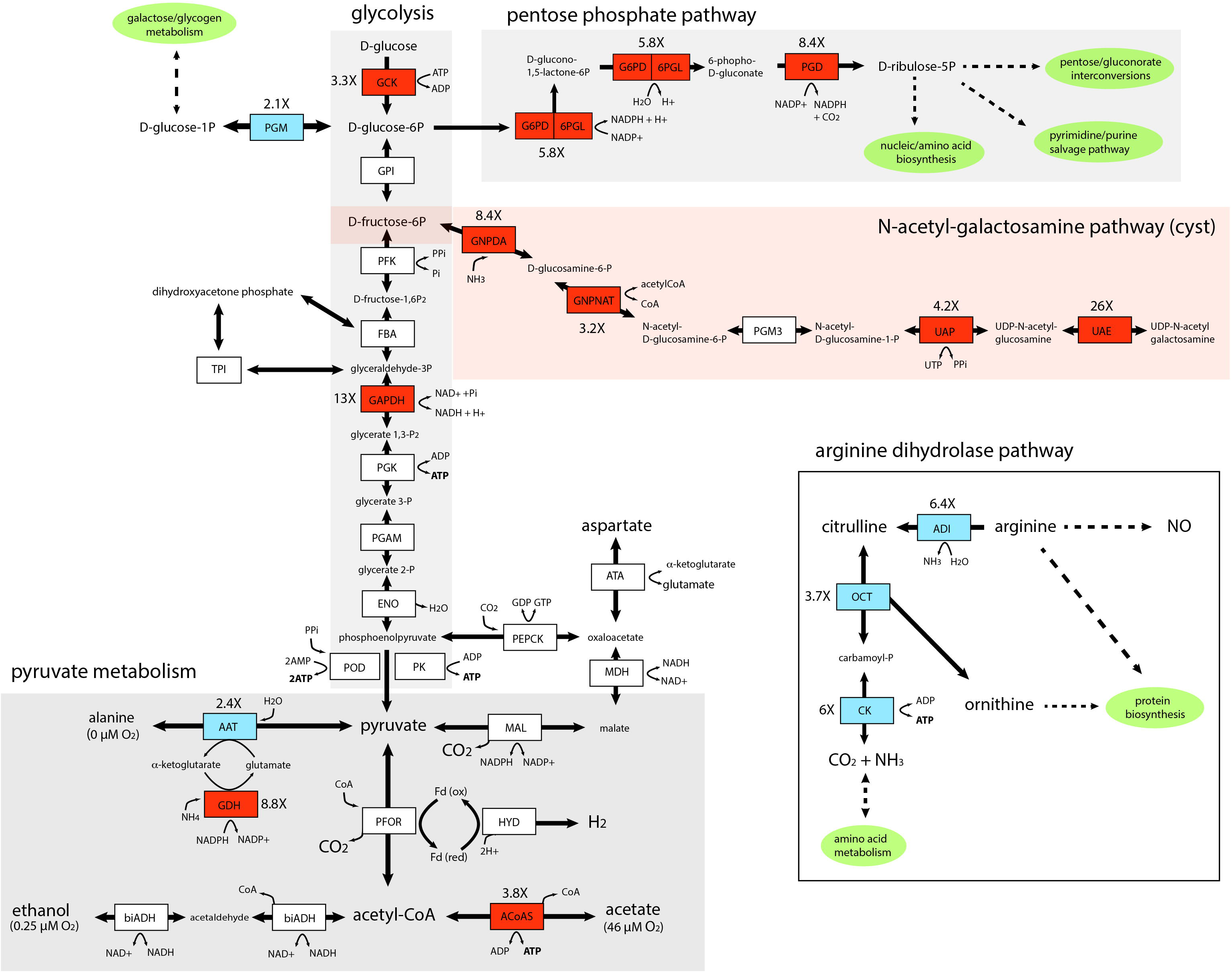
Key enzymes in the glycolytic, pentose phosphate, pyruvate, and UDP-N-acetyl-galactosamine pathways are upregulated in the *in vivo Giardia* foci. Diagrammatic representation of differentially expressed enzymes associated with *in vivo Giardia* energy and biosynthetic pathways. Red shading denotes increased expression of *in vivo* relative to *in vitro* transcripts, and blue shading denotes decreased expression. White shading indicates no significant differential expression between *in vivo* foci and *in vitro* (TYDK) transcriptomes. Oxygen concentrations associated with different branches of the pyruvate metabolic pathway are noted [56]. The enzyme abbreviations, names, GiardiaDB (50803) ORFIDs, and Enzyme Commission numbers are: <u>Glycolysis</u>: ACYP. acylphosphatase (7871), EC 3.6.1.7; ENO, enolase (11118), EC 4.2.1.11; FBA, fructose-bisphosphate aldolase (11043), EC 4.1.2.13; GAPDH, glyceraldehyde-3-phosphate dehydrogenase (17043), EC 1.2.1.12; GPI, glucose-6-phosphate isomerase (9115), EC 5.3.1.9; GCK, glucokinase (8826), EC 2.7.1.2; PFK, phosphofructokinase (14993), EC 2.7.1.90; PGAM, phosphoglycerate mutase (8822), EC 5.4.2.1; PGK, phosphoglycerate kinase (90872), EC 2.7.2.3; PGM, phosphoglucomutase (17254), EC 5.4.2.2; PK, pyruvate kinase (17143,3206), EC 2.7.1.40; POD, pyruvate:orthophosphate dikinase (9909), EC 2.7.9.1; TPI, triose phosphate isomerase (93938), EC 5.3.1.1. <u>Pyruvate metabolism</u>: ATA, aspartate transaminase (91056), EC 2.6.1.1; FeADH, Fe-alcohol dehydrogenase (3861, 3593) EC 1.1.1.1; ACoAS, acetyl-CoA synthetase (13608); AAT, alanine aminotransferase (12150, 16353), EC 2.6.1.2;biADH (bifunctional alcohol/aldehyde dehydrogenase E (93358), EC 1.1.1.1; GDH, NADP-specific glutamate dehydrogenase (21942) EC 1.4.1.3, 1.4.1.4; Fd, ferredoxin (9662, 27266,10329); HYD, hydrogenase (6304), EC 1.12.7.2; MAL, malic enzyme (14285), EC 1.1.1.38; MDH, malate dehydrogenase (3331), EC 1.1.1.37; PEPCK, phosphoenolpyruvate carboxykinase (10623), EC 4.1.1.32; PFOR, pyruvate:ferredoxin oxidoreductase (17063, 114609); <u>Pentose Phosphate Pathway (PPP)</u>: G6PD-6PGL, glucose-6-phosphate-1-dehydrogenase (8682), EC 1.1.1.49; PGD, phosphogluconate dehydrogenase (14759), EC 1.1.1.44; <u>UDP-N-acetylgalactosamine (GalNac) biosynthetic pathway</u>: GNPDA, glucosamine-6-phosphate deaminase (8245), EC 3.5.99.6; GNPNAT, glucosamine 6-phosphate *N*-acetyltransferase (14651), EC 2.3.1.4; PGM3, phosphoacetylglucosamine mutase (16069), EC 5.4.2.10; UAE, UDP-*N*-acetylglucosamine 4-epimerase (7982), EC 5.1.3.2; UAP, UDP-*N*-acetylglucosamine diphosphorylase (16217), EC 2.7.7.23. <u>Arginine Dihydrolase Pathway</u>: ADI, arginine deaminase (112103), EC 3.5.3.6; ARG-S, arginyl-tRNA synthetase (10521), EC 6.1.1.19; CK, carbamate kinase (16453), EC 2.7.2.2; NOS, nitric oxide synthase (91252), EC 1.14.13.39; OCD, ornithine cyclodeaminase (2452), EC 4.3.1.12; OCT, ornithine carbamoyltransferase (10311), EC 2.1.3.3; ODC, ornithine decarboxylase (94582), EC4.1.1.17; PRO-S, prolyl-tRNAsynthetase (15983), EC 6.1.1.15;

The carbohydrate component of the cyst is comprised of a D-GalNAc(b1,3)-D-GalNAc homopolymer synthesized from fructose 6-phosphate via the N-acetylgalactosamine (GalNac) biosynthetic pathway (Figure 4). As has been seen previously during the transcriptional activation of GalNAc pathway enzymes during *in vitro* encystation [54; 55], we found that almost all enzymes in this pathway were expressed between 3.2 to 26X higher in the *in vivo* foci as compared to in the *in vitro* transcriptomes. In fact, the second enzyme in this pathway, glucosamine 6-phosphate *N*-acetyltransferase (GL50803_14651), is one of the top 25 highly expressed genes in *in vivo* foci (Figure 3B and Supp table).

Key enzymes involved in the generation of ATP via glycolysis (GCK, 3.3X, and GAPDH, 13X) and pyruvate metabolism (AcoAS, 3.8X) were also more highly transcribed in the foci. PGM expression was several fold (2.1X) lower in the foci transcriptome relative to *in vitro* transcriptome. Pyruvate metabolism in *Giardia* is sensitive to oxygen concentration; at lower oxygen concentrations, pyruvate is fermented to alanine and ethanol, whereas at higher oxygen concentrations pyruvate is converted to acetate with the concomitant production of ATP [56]. In foci, we observed increased expression of AcoAS, a key enzyme in the higher oxygen pathway resulting in acetate production. We also noted that AAT, involved in alanine production, was less expressed in foci than *in vitro* culture (-2.4X). Lastly, two key enzymes of the pentose phosphate pathway (G6PD, 5.8X and PGD, 8.4X) are more highly expressed in the *in vivo* foci transcriptomes (Figure 4).

### Partitioning analysis identifies highly expressed in vivo foci genes associated with encystation

Partitioning analysis sorted nine gene clusters of similar expression levels, and each cluster ranged from between 39 and 218 differentially expressed genes. On average, about 50 percent of the genes identified using the differential expression analyses (Methods) were shared with genes in clusters identified using the partitioning analysis (Figure 5B). The gene clusters that were the most highly transcribed in the foci (cluster A, 15X, cluster B, 5X and cluster C, 3X, respectively) were enriched for encystation, oxidative stress response, and membrane transporter genes (Figure 5C). Cluster A included the most encystation-associated genes (Figure 5B and Supplemental Tables 7 and 8). These three clusters also shared genes in common with several recent studies of *in vitro* encystation genes (Morf et al.), putative cyst-specific genes (Palm et al.), and putative host-induced parasite genes (Ringqvist et al.) (Figure 5B).

**Figure 5.**
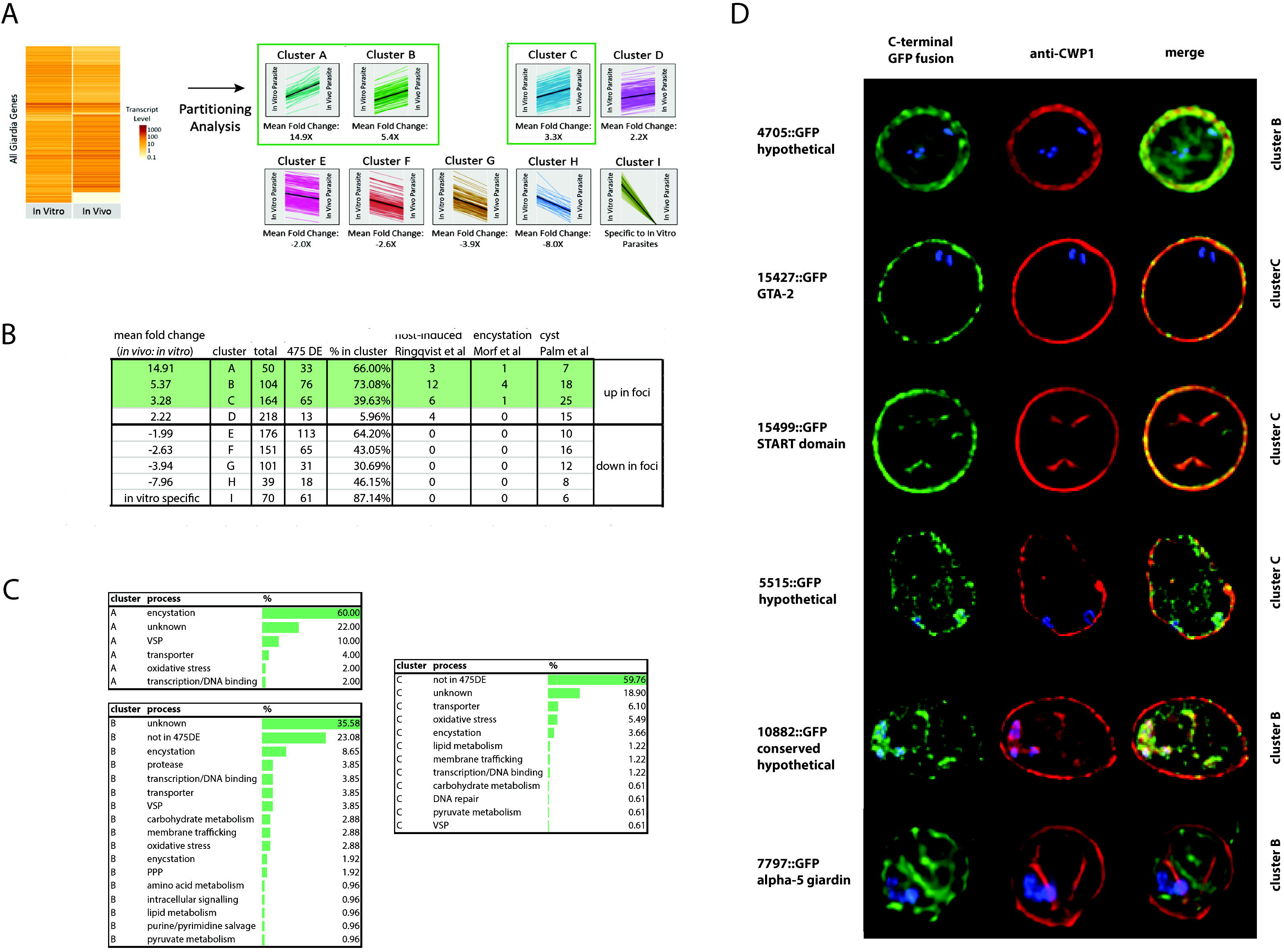
Identification and confirmation of novel proteins expressed highly *in vivo* that localize to cysts. Differentially expressed genes from the *in vivo* foci and *in vitro* culture datasets were partitioned into nine gene clusters (A-I) based on JS distance (Methods). Many of the 475 differentially expressed genes classified using both CUFFDIFF and SLEUTH (Methods) are highly represented in the clusters from this study (B). Other host-induced or encystation-specific transcriptome studies (Ringqvist et al [113], Morf et al [34], Palm et al [50]) have fewer genes represented in the clusters than those identified by partitioning analysis in the foci (B). Many genes in clusters A, B, and C are associated with encystation or oxidative stress (C). In D, C-terminal GFP fusions of genes from cluster B and C also are associated with the cyst wall or the interior of the cyst as visualized using colocalization immunofluorescent images with the cyst wall protein 1 (CWP-1) antibody.

### Confirmation of six new cyst-associated proteins expressed at higher levels in the in vivo foci

To evaluate whether some of the genes of unknown function identified using either differential expression or partitioning analyses were associated with encystation, we tagged fifteen genes using C-terminal GFP tags and created GFP-fusion strains [51]. Based on fluorescence microscopy of trophozoites of these GFP fusion strains, thirteen of the fifteen strains had interphase localization in trophozoites (Supplemental Figure 1). Six had localization in cysts (Figure 5D), with three localizing to the cyst wall: GTA-2 (15427), a START domain protein (15499), and one hypothetical protein (4705). Three GFP fusions localized to the interior region of the mature cyst including alpha-5-giardin (7797), a conserved hypothetical protein (10882), and one hypothetical protein (5515). The 4705::GFP and 7797::GFP strains lacked any localization in the trophozoite stage. Fifteen other genes more highly expressed *in vitro* culture were also tagged, and fourteen had localizations in trophozoites (Supplemental Figure 2).

## Discussion

Defining the *in vivo* dynamics of encystation and cyst dissemination is key to understanding *in vivo* host-parasite interactions and *Giardia’s* pathogenesis [34; 57; 58]. The *Giardia* cyst is characterized by four nuclei and a thick outer cell wall consisting of cyst wall proteins 1, 2, and 3 and a unique β-1,3-linked N-acetylgalactosamine homopolymer. The structure of the cyst wall allows cysts to persist in the environment until they are ingested by a compatible host [14; 15; 59; 60; 61]. Encystation has been proposed to be triggered by differential environmental chemistry associated with distinct regions of the gastrointestinal tract (i.e. the distal small intestine and colon), and has thus long been assumed to occur when trophozoites are swept towards the large intestine in mammalian hosts [14; 15; 59]. *In vitro* culture conditions for encystation have been developed that are thought to mimic the high pH and limiting cholesterol that trophozoites encounter in the gut [62]. While landmark *in vitro* studies established that the initiation of encystation is transcriptionally controlled [8; 9; 10], our understanding of the spatiotemporal dynamics of parasite metabolism and differentiation in the host is limited due to the inherent challenges of evaluating *Giardia* physiology *in situ* in the gastrointestinal tract.

Recent work in our laboratory challenges the paradigm that encystation is triggered late in infection via spatially segregated cues in the gastrointestinal tract [40]. By directly imaging *Giardia* infections in the mouse using *in vivo* and *ex vivo* bioluminescent imaging (BLI) of parasites with integrated firefly luciferase bioreporters of metabolism (*P_GDH_-Fluc* strain) or encystation (*P_CWP1_-Fluc* and *P_CWP2_-Fluc* strains), we showed that parasites colonize both the proximal and distal small intestine in high-density foci early in infection. Encystation is also initiated early during infection in these foci, and persists throughout the entire course of infection. Barash et al. also confirmed that trophozoites extracted from proximal small intestines of infected mice displayed encystation-specific vesicles (ESVs), which develop during the differentiation of the trophozoite to the cyst [40]. This work predicted that that trophozoite colonization and encystation occur simultaneously in the host, and showed that encysting trophozoites were enriched in the high-density foci.

Here we used the same BLI *ex vivo* imaging strategy to sample high-density *Giardia* foci in the proximal small intestine of four animals infected with a constitutive luciferase strain *P_GDH_-Fluc* (Figure 1 and [40]). Despite our deriving the samples from the gastrointestinal tracts of four different animals either three or seven days post *Giardia* infection, the transcriptomic profiles of the *in vivo* foci were strikingly similar (Figure 2A). Thus, we predict that trophozoites synchronize their physiological responses once they reach a threshold density in regions of the proximal small intestine and are committed to encystation (Figure 2).

### *In vivo* Giardia transcriptional profiles are dominated by encystation

Using two different analytic methods (Figure 2B and 2C), we found that the transcriptome profiles of *Giardia* from *in vivo* foci and log phase axenic cultures were significantly different. Relative to *in vitro* culture, many of the most highly expressed genes in the *in vivo* foci have functions associated with both encystation and oxidative stress response. In contrast, the top *in vitro* expressed genes are associated with translation, cell division, and arginine metabolism as has been previously reported (Figure 2D and 2E and [63; 64]).

We have previously shown that the CWP1 and CWP2 promoter fusions to luciferase are highly expressed in the *in vivo* foci using BLI, and confirmed that trophozoites in high-density foci express numerous encystation specific vesicles (ESVs) [40]. The biogenesis of ESVs that transport cyst wall proteins to the plasma membrane of the trophozoite is a hallmark of encystation. ESVs are synthesized in the endoplasmic reticulum and transported to the periphery of the cell in late encystation. Cyst wall proteins are sorted and transported by novel large vesicles, encystation secretory vesicles (ESVs) and ultimately by peripheral vesicles to the growing cyst wall. Many other components of the cyst have also been suggested by expression or proteomic analyses or confirmed by localization, including protein disulfide isomerases [65], high cysteine membrane proteins (HCMPs) [66], and EGFP family proteins [66].

In support of our assertion that *Giardia* from *in vivo* foci are already committed to encystation [40], we demonstrate that the transcriptome of *in vivo* foci is defined by the high expression of a number of genes associated with cyst wall biosynthesis. Several aspects of trophozoite differentiation to cysts are transcriptionally regulated [60]. The leucine-rich repeat proteins, cyst wall proteins (CWP1-3) and the enzymes of the GalNAc biosynthetic pathway are all transcriptionally upregulated by the transcription factor Myb2 within 90 minutes of switching trophozoite cultures to encystation medium [67]. GalNAc enzyme transcription has been shown to peak at about 22 hours in encystation medium [39]. In the *in vivo* foci, we noted significantly higher expression of genes associated with encystation, including the transcriptional regulator Myb2, all enzymes of the GalNAc biosynthetic pathway (Figure 4), two known cyst wall proteins (CWP2 and CWP3), 17 HCMPs, and two EGF family proteins [68] (Figures 2, 3, and Supplemental Table 4). CWP1 [69] was identified as differentially expressed by only one of the two methods of analysis we used, and was thus not included in the 475 differentially expressed genes (Methods). Through partitioning analysis and analysis of differential gene expression in the *in vivo* foci, we also identified six new cyst-specific proteins (Figure 5D and Supplemental Tables 7 and 8). Each localizes to the cyst wall or the cyst interior after 24 hours of incubation in encystation medium.

Variant-specific surface proteins (VSPs) are cysteine-rich membrane proteins of variable size that are responsible for antigenic variation and likely contribute to *Giardia’s* evasion from the host immune response [70]. The *Giardia* genome encodes over seventy variant surface protein (VSP) genes, yet only one VSP is expressed per cell [71]. VSP expression occurs early during *in vitro* encystation [72], and there are dramatic shifts in expression of VSP genes late in *in vitro* encystation [39]. Expressed VSPs are then transported by ESVs during encystation [6]. Fourteen VSPs were differentially expressed in our analysis, and two VSPs were among the most highly expressed proteins in *in vivo* foci relative to *in vitro* culture (Figure 2).

A common symptom of giardiasis is fat malabsorption, with markedly increased lipid concentrations in luminal contents and feces [6; 73; 74]. In addition to suffering from steatorrhea, patients excreting *Giardia* cysts have altered lipid profiles in circulating blood as well as in the gut lumen [75]. *Giardia* directly perturbs lipid metabolism within the host by scavenging the gut lumen for lipids and by excreting novel end products of lipid metabolism [76]; it also contributes to shifts in the local gut microbiota [12] by both excreting novel lipids and influencing the bioavailability of bile acids [73; 77; 78; 79].

Lipids also play key roles in the regulation of encystation [76]. Glucosylceramide transferase-1 (gGlcT1), an enzyme involved in sphingolipid biosynthesis that has also been shown to play a key role in ESV biogenesis and cyst viability [80], is highly expressed in *in vivo* foci. Also highly expressed in foci is acid sphingomyelinase-like phosphodiesterase 3b precursor gSmase 3B. GSMase 3B has been shown to be transcriptionally upregulated during encystation and is proposed to scavenge ceramide from dietary components in the small intestine [80]. We also noted the increased *in vivo* expression of seven genes associated with membrane trafficking that could be involved in ESV biogenesis or transport (Figure 3 and Supplemental Table 4).

Five cysteine proteases (three cathepsin B homologs: 10217, 17516, 16779 and two cathepsin L homologs: 11209, 137680) have roughly between five and 11.5-fold higher expression in the foci (Supplemental Table 2 and 4). Cathepsin B (16779, CP3) and cathepsin L (137680) are known to be upregulated in encysting trophozoites [81]. In *Giardia,* cathepsin cysteine proteases are linked to not only to encystation [82] but also to evasion of innate and adaptive immune responses. In co-culture with rat IEC epithelial cells, one (3099) of the nine cathepsin B proteases genes has significantly increased expression *Giardia* trophozoites [83]. Cathepsin B proteases secreted by trophozoites are sufficient to prevent inflammatory responses from the host by its degradation of the chemokine CXCL8, inhibiting chemotaxis of pro-inflammatory PMN cells to the site of parasite colonization [84].

In total, we have confirmed the expression of 37 known encystation genes and identified six new cyst-associated proteins in the high-density foci. Because we have identified many highly expressed encystation-associated transcripts in key metabolic, biosynthetic, protease, and membrane trafficking pathways, we suggest that the majority of cells in the high-density foci in the proximal intestine are in mid-to late stages encystation (Figure 6 and Supplemental Table 9). The apparent developmental synchrony of cells in the foci is supported by the strong encystation transcriptional profiles in each sample regardless of the day of infection (Figure 6) or in each animal sampled (Figure 2A) and may be the result of density related cues such as nutrient or lipid deprivation that trigger encystation.

**Figure 6.**
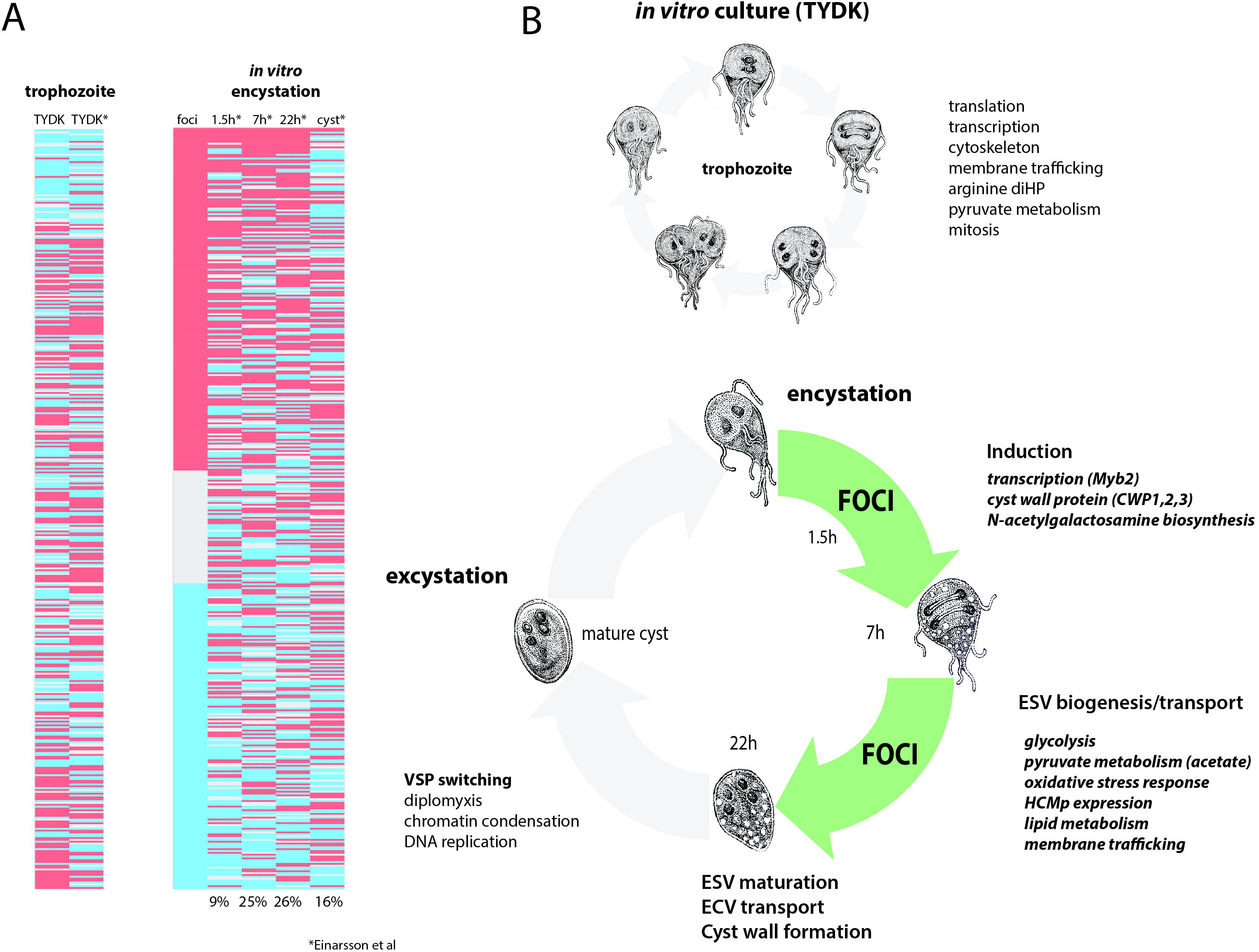
Differentially expressed genes in *in vivo* foci are similar to mid-late encystation time points in *in vitro* encystation transcriptomes. *In vitro* and *in vivo* differentially expressed genes from this study are compared to *in vitro* encystation transcriptome of Einarsson et al. Panel A compares the 475 differentially expressed genes in this study (red = upregulated in foci; blue = downregulated in foci) to *in vitro* culture transcriptome (TYDK) and the transcriptomes of four time points of an *in vitro* encystation transcriptome study. Percentages indicate the number of similarly differentially expressed genes in the *in vivo* foci relative to that *in vitro* encystation time point. Panel B summarizes processes upregulated in *in vitro* culture, which is primarily trophozoite growth and division (above) as contrasted with cellular processes upregulated in the *in vivo* foci (green) that are more similar to those genes upregulated during in mid-to-late *in vitro* encystation.

### Energy metabolism in the *in vivo* foci

As an anaerobic fermentative microbe, *Giardia* uses glycolysis, pyruvate fermentation and the arginine dihydrolase pathways for energy production ([2] and see Figure 4). Though trophozoites cannot tolerate high oxygen concentrations, they actively consume oxygen [85; 86] and may take advantage of slightly elevated oxygen levels for energetic benefit [6; 63; 87]. Thus, glucose is fermented to alanine, ethanol, acetate and CO2, but the end product balance of parasite fermentation varies with local oxygen tension and glucose levels, with the highest NAD(P)+ or ATP yields occurring at the higher end of tolerated oxygen concentrations [63; 88; 89; 90]. Under strictly anaerobic conditions, pyruvate is converted into alanine and NADP is regenerated. At very low oxygen concentrations (less than 0.25 μΜ), alanine production is inhibited and pyruvate is converted primarily to ethanol with the regeneration of NAD [10]. As oxygen levels increase above 0.25 μM, the ATP producing arm of pyruvate metabolism is activated and acetate and ATP become the primary end products. At these higher oxygen concentrations, it is hypothesized that antioxidant enzymes consume NAD(P)H to neutralize dO2 and ROS/RNS, leading to regeneration of NAD(P) for glycolysis, allowing pyruvate to be used for ATP production, instead of NAD(P)H regeneration [10].

We noted higher expression of genes associated with energy metabolism via glycolysis (GCK, 3.3X and GAPDH, 13X) and pyruvate fermentation via pathways associated with higher oxygen tensions (Figure 4). These expression patterns are indicative of redox stress and encystation-specific metabolism associated with the *in vivo* foci. *In vivo,* acetate could be the primary end product of *Giardia* fermentation, rather than ethanol or alanine, due to the relatively increased expression of ACoAS (3.8X) and the decreased expression of AAT (2.4X) we observed (Figure 4). We predict the increased expression of GDH is associated with parasite ammonia assimilation and the cycling of NAD(P) to maintain intracellular redox balance [91].

Encysting trophozoites slow their catabolism of glucose for energy and switch to using glucose to synthesize N-acetylgalactosamine for cyst wall synthesis ([55] and Figure 4). GalNAc is synthesized only during encystation via a transcriptionally induced pathway of five enzymes that produces UDP-GalNac from fructose-6-phosphate that is diverted from ATP production in glycolysis to cyst wall polysaccharide biogenesis. This diversion could result in a net loss of ATP synthesis during encystation.

While little is known about the regulation of glycolysis and energy production during encystation, it is thought that additional ATP is generated from the arginine dihydrolase pathway (ADiHP) (Figure 4). This pathway yields ATP via the conversion of arginine to ammonia and citrulline, with the substrate-level phosphorylation of citrulline yielding ornithine and carbamoylphosphate, and resulting in NH_3_, CO_2_ as end products [90]. Because *Giardia* and epithelial cells use arginine for growth, the parasite and host are proposed to influence the severe malnutrition and increased susceptibility to villus shortening observed during *Giardia* infection [92; 93]. Ornithine carbamoyltransferase (OCT) is one of the most highly expressed enzymes in culture and in the *in vivo* foci, yet we observed less expression of key enzymes in the ADiHP relative to *in vitro* culture (Figure 4). This is counter to the prevailing notion that encysting trophozoites increase arginine catabolism through the ADiHP to offset energy lost from slowing glycolysis [64].

Lastly, trophozoites also lack purine and pyrimidine synthesis, relying solely on salvage pathways to obtain these metabolites from the host and commensal microbiota [94; 95; 96]. We also discovered that several key enzymes of the pentose phosphate pathway (G6PD 5.8X and PGD 8.4X) have increased expression in the *in vivo* foci transcriptome (Figure 4). A key product of the pentose phosphate pathway (PPP) is NADPH, which plays an integral role in reductive biosynthesis as well as in the defense against oxidative stress and the maintenance of cellular redox. In *Giardia* and in *Trichomonas,* the G6PD and 6PGL enzymes are fused and are directly involved in the oxidative branch of the PPP in generating reduced NADPH (Figure 4 and [97]). This suggests that in the foci, there is increased cycling of reducing equivalents with CO2 production and the increased production of D-ribose-5P as a key substrate for nucleotide and amino acid biosynthesis and pyrimidine/purine salvage pathways.

### Oxidative stress responses are increased in the *in vivo* foci

Dynamic changes in the gut oxygen concentrations affect the formation of nitric oxide and reactive oxygen species (ROS). *Giardia* attaches to the villi of the small intestine, where oxygen levels range up to 60 μΜ [6]. Anti-oxidant defense is thought to be mediated by the thioredoxin-thioredoxin reductase system, as *Giardia* lacks superoxide dismutase, catalase, and the glutathione-glutathione reductase. This system includes an FAD containing NADPH-dependent disulfide reductase, a thioredoxin, a thioredoxin reductase (TrxR) and a thioredoxin-peroxidase [98]. As *Giardia* cysts do consume oxygen [86], the thioredoxin-thioredoxin reductase system may play a role in the maintenance of oxidation/reduction homeostasis in both trophozoites and cysts.

*Giardia* gene expression is modulated by oxygen and other stressors, and a microaerophilic environment may improve *Giardia’s* energy metabolism. Oxygen depletion thus may reduce the efficiency of glycolysis *in vitro,* and byproducts from antioxidant enzymes are proposed in *Giardia* to promote glycolysis and energy production [6]. In oxidative environments, *Giardia* expresses NADH oxidase and flavodiiron protein that metabolize oxygen to form water [99; 100]. Nitrosative stresses can have differing metabolic effects, however; while high micromolar concentrations of nitric oxide are toxic [101], lower levels inhibit cell proliferation, encystation, and excystation [102]. Resolution of nitric oxide stress by *Giardia* is believed to be mediated by activity of its flavohemoglobin protein [103], which catalyzes the oxidation of nitric oxide to nitrate using oxygen as a co-substrate [104; 105; 106; 107].

Transcriptomic analyses of the high-density foci suggest that they are susceptible to and are responding to localized oxidative and nitrosative stresses [99; 100]. In particular, we observed the increased expression of genes associated with oxidative or nitrosative stress and detoxification, including NADPH oxidoreductase (17150, 15004), nitroreductase (FMN) family protein (8377), and flavohemoglobin (15009) at levels that were 2.5-8.5 fold higher than *in vitro* culture. In pathogenic bacteria such as *E. coli* and *Salmonella,* flavohemoglobin activity is significantly increased *in vivo* intestinal infections, coincident with host nitric oxide production [108; 109]. In response to nitrosative stress (in the form of exposure to >5 mM nitrite), *in vitro* cultures displayed enhanced flavohemoglobin expression, and trophozoites exhibited more efficient metabolism of nitric oxide [110]. Indeed many of the top 25 expressed genes in the *in vivo* foci (e.g. NADPH oxidoreductase) are associated with oxidative or nitrosative stress responses [111], which may explain how *Giardia* can colonize the proximal part of the small intestine, a fairly aerobic portion of the intestinal tract [110].

Thioredoxin associated proteins also aid in the detoxification of oxidative stress, and five thioredoxin domain proteins (9827, 6289, 2388, 8064, 9335) were also highly expressed in the *in vivo* foci (Figure 2 and 3). One thioredoxin reductase (TrxR, 9827) has disulfide reductase activity and converts oxidized thioredoxin to its reduced form. Reduced thioredoxin is, in turn, used to prevent protein misfolding that can occur under oxidative stress conditions [8]. *Giardia’s* TrxR also has NADPH oxidase activity, and during conversion of thioredoxin to its reduced form, NADP is generated from NADPH [16].

Due to these transcriptomic patterns, we predict that *Giardia* trophozoites in high-density *in vivo* foci are more likely to be subject to host-induced oxidative stressors than *Giardia* growing at lower density. Parasites in the high-density foci may take advantage of a more oxidized environment to switch to more energetically favorable pyruvate fermentation.

### Density dependence of encystation and the consequences of encystation on host pathology

The physiology of parasites within the high-density foci in the host gut clearly differs from that of cells in laboratory culture or in co-culture with cell lines. The observed encystation-and oxidative stress-specific responses in gene expression within parasite foci may reflect physiological changes associated with high-density growth in localized regions of the gut. We recently showed that increased density in culture results in the upregulation of encystation-specific promoters in *Giardia* [40]. Density-associated differentiation also occurs other protists, such as the amoeba *Dictyostelium discoideum,* wherein the accumulation of a factor secreted by starved cells leads to initiation of multicellular aggregation and assembly of a fruiting body [17]. Differentiation of *Trypanosoma brucei* is also dependent on cell density. When the slender, proliferative bloodstream form reaches a specific threshold density, accumulation of secreted ‘stumpy induction factor’ induces differentiation to the stumpy, transmissive form that is taken up by the tsetse fly [18]. At high-density in the mouse, *Giardia* responds in a way that promotes differentiation and transmission.

The similarity of transcriptional profiles between all sampled foci indicates that trophozoites have already committed to encystation, suggesting they have received the inducing encystation stimulus within the last 12 hours [58]. We hypothesize that once parasites reach threshold high densities in discrete regions of the gut, they could trigger localized host immune responses and redox shifts, resulting in the observed increase in oxidative stress and encystation responses that define transcription in the foci. Current hypotheses about how giardiasis induces host symptoms have focused primarily on specific damage induced by parasite attachment to the host gut epithelium or on the production of anti-microbial metabolites by the mammalian host.

Lastly, we have recently reported that *Giardia* infection results in a dysbiosis throughout the gastrointestinal tract in mice, characterized by a shift in the diversity of commensal microbiota towards more aerobic Proteobacteria species [112]. As we find that encystation-specific metabolism occurs early and consistently during *Giardia* infection in the host, it is likely that nutrient deprivation and waste production during encystation would affect the nutrition of the host and associated commensal microbiome. Further investigation of Giardia-host-microbiome interactions should thus emphasize the study of such interactions *in situ* in order to vet laboratory culture and co-culture studies.

## ACKNOWLEDGEMENTS

Bioluminescent imaging was performed at the Center for Molecular and Genomic Imaging (CMGI), University of California, Davis with gracious help and training from Jennifer Fung and Charles Smith. The authors especially acknowledge Sarah Guest for the *Giardia* life cycle drawings. This study was supported by NIH R01AI077571 and NIH R21 AI119791 awards to SCD.

**Supplemental Figures**

Supplemental Figure 1: Subcellular localization of C-terminal GFP fusion proteins of selected genes with increased expression in the *in vivo* foci

Supplemental Figure 2: Subcellular localization of C-terminal GFP fusion proteins of selected genes with increased expression in *in vitro* axenic culture

**Supplemental Table Legends**

Supplemental Table 1: Quantification of whole body and *ex vivo* bioluminescent imagine at day 3 and day 7 p.i. for transcriptomic sampling of foci

Supplemental Table 2 FPKM of *in vivo* and *in vitro* transcriptomes

Supplemental Table 3 Overall heatmap of all FPKM

Supplemental Table 4 Heat map table unregulated

Supplemental Table 5 Heat map table - down regulated

Supplemental Table 6 Heat map table - *in vitro* only

Supplemental Table 7 Cuffdiff partitioning analysis into nine clusters

Supplemental Table 8 Annotated nine clusters

Supplemental Table 9 Comparison of *in vivo* foci and *in vitro* encystation transcriptomes

